# Variations in Brain Morphometry Among Healthy Preschoolers Born Very Preterm

**DOI:** 10.1101/734277

**Authors:** Holly M. Hasler, Timothy T. Brown, Natacha Akshoomoff

## Abstract

**Background:** Preterm birth is associated with an increased risk of neonatal brain injury, which can lead to alterations in brain maturation. Advances in neonatal care have dramatically reduced the incidence of the most significant medical consequences of preterm birth. Relatively healthy preterm infants remain at increased risk for subtle injuries that impact future neurodevelopmental and functioning.

**Aims:** To investigate the gray matter morphometry measures of cortical thickness, surface area, and sulcal depth in the brain using magnetic resonance imaging at 5 years of age in healthy children born very preterm.

**Study design:** Cohort study

**Subjects:** Participants were 52 children born very preterm (VPT; less than 33 weeks gestational age) and 37 children born full term.

**Outcome measures:** Cortical segmentation and calculation of morphometry measures were completed using FreeSurfer version 5.3.0 and compared between groups using voxel-wise, surface-based analyses.

**Results:** The VPT group had a significantly thinner cortex in temporal and parietal regions as well as thicker gray matter in the occipital and inferior frontal regions. Reduced surface area was found in the fusiform area in the VPT group. Sulcal depth was also lower in the VPT group within the posterior parietal and inferior temporal regions and greater sulcal depth was found in the middle temporal and medial parietal regions. Results in some of these regions were correlated with gestational age at birth in the VPT group.

**Conclusions:** The most widespread differences between the VPT and FT groups were found in cortical thickness. These findings may represent a combination of delayed maturation and permanent alterations caused by the perinatal processes associated with very preterm birth.

## Variations in Brain Morphometry Among Healthy Preschoolers Born Very Preterm

Preterm birth is associated with an increased incidence of brain injury at birth that involves primarily the periventricular white matter and subcortical structures. This has been termed the “encephalopathy of prematurity” and is the primary determinant of neurodevelopmental difficulties in childhood (Volpe, 2009). Despite dramatic advances in neonatal care and increased survivability, individuals born very preterm (VPT), at less than 33 weeks gestation, continue to show differences in development of brain structures throughout childhood and as adults. These differences may be attributed to complex processes that involve brain injury at birth as well as longer term neurodevelopmental differences (Ortinau & Neil, 2015). Early injury has been shown to affect proximal and distal structures as the brain develops into childhood and beyond. The pattern of these complex findings suggests permanent cortical changes in some regions (perhaps due to neuronal loss in the perinatal period) and delayed development of later maturing cortical regions and white matter tracts. Preschool age is a time of rapid development of the brain, characterized by expansion of cortical surface area and contraction of gray matter thickness (Brown et al., 2012). Little is known about how these processes are affected in preschool age children who were born VPT.

Previous studies have demonstrated changes in cortical morphometry as a consequence of preterm birth. These studies have demonstrated some areas where thickness is greater in individuals born VPT, and other areas with lower thickness compared to individuals born full term (FT). Cortical thickness was lower in middle temporal, anterior cingulate, and posterior inferior parietal regions during middle childhood and adolescence (Lax et al., 2013; Martinussen et al., 2005; Nagy, Lagercrantz, & Hutton, 2011). Other areas have been found to have greater thickness in VPT children, adolescents, and young adults, primarily within the temporal pole and medial and lateral frontal brain regions (Martinussen et al., 2005; Mürner-Lavanchy et al., 2014; Nam et al., 2015) as well as medial occipital and anterior cingulate regions (Lean, Melzer, Bora, Watts, & Woodward, 2017).

Previous studies have found mixed results regarding the effect of preterm birth on measures of cortical surface area. In school-age children born with very low birthweight (VLBW), smaller surface area was found in the bilateral medial and lateral temporal lobes, inferior frontal lobes, and parietal/occipital regions (Sripada et al., 2018). Reduced surface area was also found in a group of 3-to 4-year-old VLBW children in orbitofrontal and transverse temporal regions (Phillips et al., 2011), as well as in studies of VLBW older children and young adults (Grunewaldt et al., 2014; Skranes et al., 2013; Solsnes et al., 2015).

Sulcal depth may also be sensitive to the effects of preterm birth. There is evidence that preterm birth may alter the timing and trajectory of the deepening of sulci (Dubois et al., 2008). In a study of cortical folding, VPT-born infants showed more shallow sulci than FT infants at term-equivalent age (Engelhardt et al., 2015). Shallower sulci have been reported in the superior temporal sulci and inferior portion of the pre and post-central sulci in children born VPT (Zhang et al., 2015).

The purpose of this study was to characterize the brain structural properties of a group of healthy preschoolers born VPT with relatively benign neonatal health history. This same sample was included in two of our previous papers (Hasler & Akshoomoff, 2019; Hasler et al., Submitted). Here we compared measures of surface area, cortical thickness, and sulcal depth between VPT and FT children at preschool age. Based on results from previous studies of older children, we hypothesized that the VPT group would have lower cortical thickness in the temporal and parietal lobes and greater thickness in the medial and lateral frontal lobes. We also predicted that areas of differences in surface area were likely to overlap with cortical thickness, with the VPT group having smaller surface area in the temporal and parietal regions. Finally, we hypothesized that the VPT group would have shallower sulci in the temporal lobe and medial occipital regions.

## Method

### Participants

The final sample was composed of 52 children born VPT and 37 children born FT. Children were enrolled in the study within 6 months of beginning kindergarten. The VPT group was recruited primarily from the follow-up program for two neonatal intensive care units in San Diego (UC San Diego and Sharp Mary Birch). The purpose of this study was to investigate the development of children born VPT without significant neonatal brain injury. Therefore, children were excluded from the study if they had a history of severe neonatal complications (i.e., Grade 3-4 intraventricular hemorrhage, cystic periventricular leukomalacia, moderate-severe ventricular dilation), known genetic abnormalities likely to affect development, and/or acquired neurological disorder unrelated to preterm birth. The children born FT were recruited via the UC San Diego Center for Human Development database of parents who consented to be contacted if their children might qualify for a study. Children born FT had no history of neurological, psychiatric, or developmental disorders. All children enrolled in the study were primarily English speaking with a Wechsler Preschool and Primary Scale of Intelligence – Fourth Edition (WPPSI-IV) (Wechsler, 2012) Full Scale IQ > 75. Additional exclusionary criteria included significant auditory or visual deficits and contraindications to MRI (e.g., pacemaker, metallic implants, recent dental procedures). The Institutional Review Board at UC San Diego approved all procedures, and each participant’s legal guardian gave written informed consent.

### Brain Imaging

Data were collected on a General Electric Discovery MR750 3.0 Tesla scanner with an 8-channel phased-array head coil at the Center for Functional MRI at UC San Diego. The full imaging protocol included: 1) a three-plane localizer; 2) a 3D T1-weighted inversion prepared RF-spoiled gradient echo scan using real-time prospective motion correction (PROMO) (White et al., 2010); 3) a 3D T2-weighted variable flip angle fast spin echo scan for detection and quantification of white matter lesions and segmentation of CSF; 4) a high angular resolution diffusion imaging (HARDI) scan with 30-diffusion directions, and integrated B0 distortion correction (DISCO). All data were inspected for quality during collection and at all stages of processing.

Data were processed at the UC San Diego Center for Multimodal Imaging and Genetics (CMIG). Structural T1-weighted images were processed using gradient nonlinearity correction. Cortical and subcortical segmentation was completed using FreeSurfer automated segmentation in version 5.3.0 (Fischl et al., 2002). The imaging protocol and data processing stream was developed specifically for data collection and analyses for studies involving young children as part of the Pediatric Imaging Neurocognition and Genetics project (PING) (see Jernigan, Brown, Hagler, et al. (2016) for full details). Briefly, non-linear transformation was used to correct for distortions caused by nonlinearity of the spatial encoding gradient fields and nonparametric nonuniform intensity normalization method was used to reduce the non-uniformity of signal intensity. The automated FreeSurfer pipeline was used for extraction of gray matter cortical thickness, cortical surface area, and sulcal depth (Fischl, Sereno, & Dale, 1999). Sulcal depth is measured as the distance from the deepest point of a sulcus to the mean height of the crown of the two adjacent gyri. Images were checked for movement or other scanner artifacts as well as errors in segmentation and registration by 2 trained experts. All images were assigned a number from 0 to 2 to quantify the amount of motion in the scan by a technician blinded to group membership. A value of 0 indicated no motion artifact in the scan, up to a value of 2 that indicated some motion artifact but image quality still within acceptable limits.

The final sample was composed of all the children enrolled in the study that were able to successfully complete the MRI scanning session and whose images were of acceptable quality. Of the 105 children originally enrolled in the study, five VPT- and one FT-born participants were unable to attempt the scanning procedure. Significant motion artifacts were present in the images of three VPT- and four FT-born children, therefore these were removed from the analyses. Additional subjects were removed due to significant errors in the FreeSurfer segmentation/reconstruction, which included two VPT- and one FT-born participant. All scans were reviewed by a neuroradiologist to inspect for brain abnormalities which led to one additional FT-born participant necessitating removal from the sample because of a brain abnormality. After these participants were removed from the sample, the final sample consisted of a total of 52 VPT and 37 FT children.

### Statistical Analyses

Statistical analyses were completed in SPSS, version 25 (Corporation, 2017). Demographic variables were compared between groups using Pearson *χ*^2^ and *t*-tests. A single socioeconomic status (SES) value was calculated for each child as a combination of parent-reported household income (4 levels) and years of maternal, or primary guardian, education (4 levels). This resulted in a value from 2 to 8 for each child; see our previous publication for full explanation of these levels (Hasler & Akshoomoff, 2019).

Surface area, sulcal depth, and cortical thickness were compared between groups using a surface-based, voxel-wise generalized linear model (GLM) with sex and age at scan as covariates. This method uses FreeSurfer tools as well as custom processing tools to calculate a voxel-wise general linear model of groups. The model is false discovery rate corrected for *p* = 0.05. The GLM results are then overlaid on a FreeSurfer standard average brain surface. Follow-up analyses utilized values derived from regions of interest (ROI) from the FreeSurfer automatic, sulcal-based parcellation (Desikan et al., 2006) corresponding to the areas of significant difference on the voxel-wise surface map. ROI values were compared between groups using ANCOVA controlling for sex and age at scan. Additional partial correlations were conducted to determine the relationship between the gray matter metrics and birth characteristic of the VPT group including GA and birth weight.

## Results

Group characteristics and mean values are presented in Table 1. The groups did not differ on sex (χ^2^ = .067, *p* = .795), age (*t* = −.447, *p* = .656), or SES (*t* = −.452, *p* = .655). As expected, there was a significant difference between groups in birthweight (*t* = −20.13, *p* < .001) and gestational age at birth (*t* = −26.75, *p* < .001). Mean WPPSI-IV Full Scale IQ was significantly different between groups (*t* = −2.36, *p* = .020), however the group mean scores were in the average range. Groups did not differ significantly on motion artifact ratings (χ^2^ = 3.72, *p* = .155).

**Table 1.**
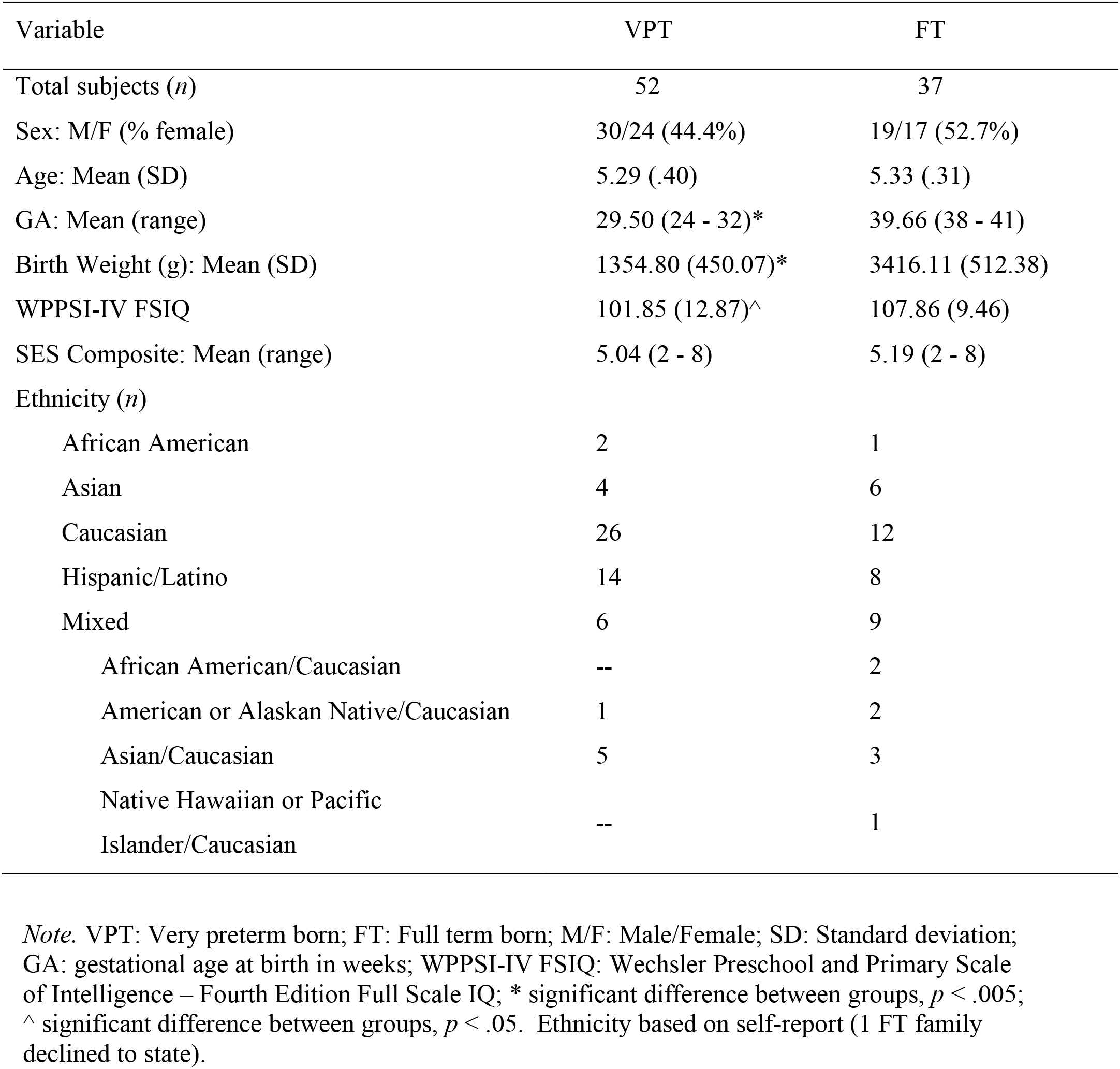
Participant Characteristics

Figure 1 depicts the differences between groups on each of the metrics of interest. Overall, the measure showing the most widespread differences between groups was cortical thickness. Children born VPT had significantly lower cortical thickness in the bilateral supramarginal and angular gyri, bilateral superior and middle temporal lobes, and bilateral superior and medial middle frontal regions. Additional areas of lower cortical thickness in the VPT group were found in the right hemisphere in lateral aspects of the frontal lobe. Children born VPT showed significantly greater cortical thickness in the cuneus and frontal pole. Total surface area was not significantly different between groups (*F*[1,85] = .081, *p* = .776). Surface area was significantly smaller in the VPT group in the fusiform gyrus. The VPT group also had significantly larger surface area in the right medial frontal, occipital pole, and anterior cingulate regions. Sulcal depth was significantly lower in the VPT group in the superior parietal lobe and medial orbital frontal regions. In summary, gray matter thickness showed the most widespread differences among the groups, with children born VPT showing significantly lower thickness in many regions of the temporal, parietal, and frontal lobes. The occipital lobe was also greatly affected in children born VPT, with significant differences across measures of thickness, and sulcal depth. Group comparisons and mean values for ROIs corresponding to regions on surface maps are displayed in Table 2.

**Figure 1.**
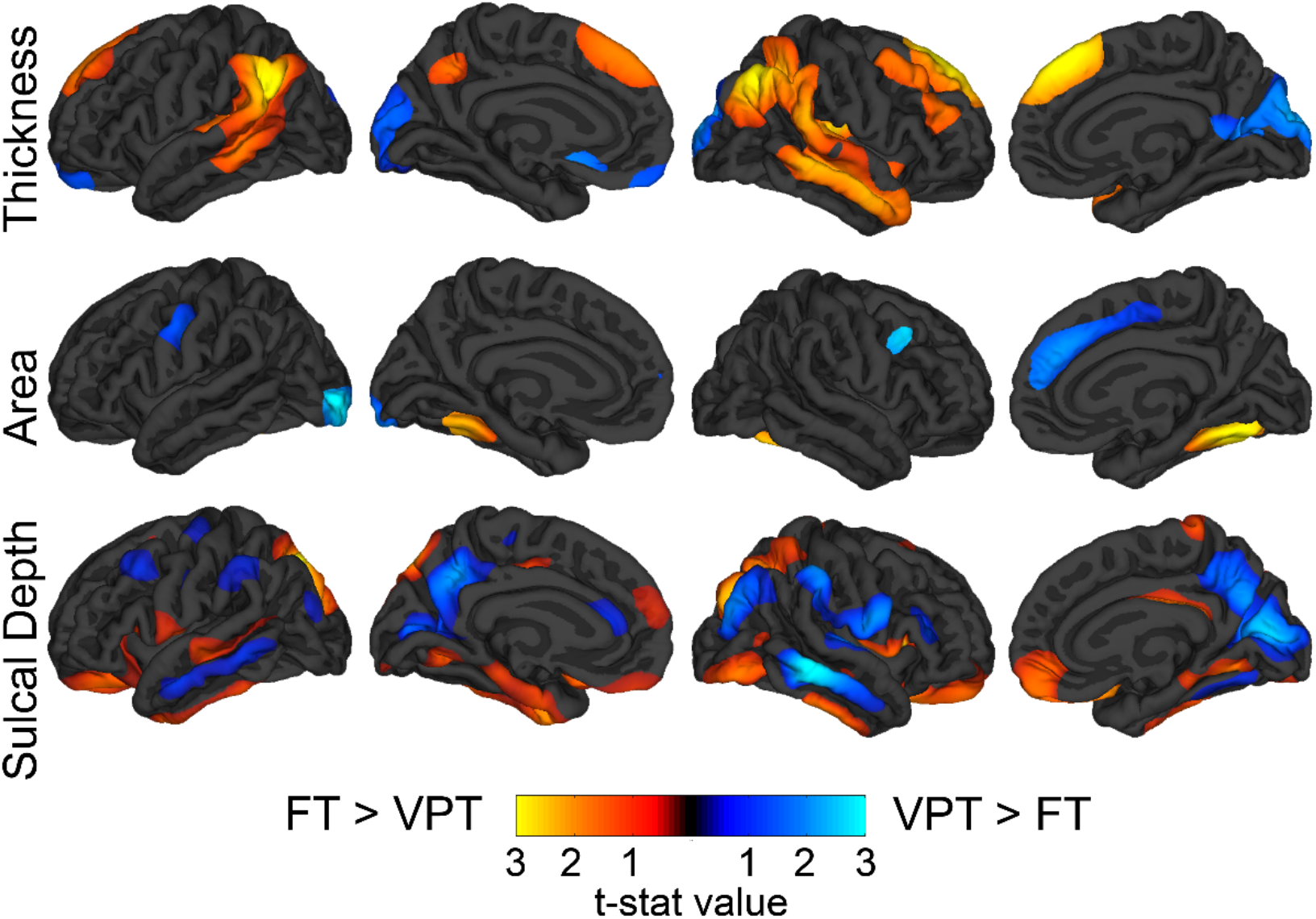
Vertex-wise comparison of gray matter measures. *Note.* Mean difference between VPT and FT groups corrected for false discovery rate = .05, projected onto the FreeSurfer average surface. Minimum threshold of *p* = .01 based on the t-statistic. FT: Full Term group; VPT: Very Preterm group

**Table 2.** ROI values for areas shown to have significant between-group differences on the vertex-wise maps

| Thickness | VPT (M, sd) | FT (M, sd) | *F* | *p* | $\eta_p^2$ |
| --- | --- | --- | --- | --- | --- |
| L inf parietal | 2.88 (.12) | 2.96 (.12) | 11.875 | .001 | .123 |
| R inf parietal | 2.94 (.13) | 3.02 (.12) | 9.502 | .003 | .101 |
| L supramarginal | 2.98 (.14) | 3.07 (.12) | 10.752 | .002 | .112 |
| R supramarginal | 3.01 (.14) | 3.08 (.12) | 6.036 | .016 | .066 |
| L middle temporal | 3.21 (.16) | 3.32 (.14) | 11.670 | .001 | .121 |
| R middle temporal | 3.25 (.13) | 3.37 (.14) | 16.547 | <.001 | .163 |
| L sup temporal | 3.18 (.14) | 3.23 (.14) | 2.713 | .103 | .031 |
| R sup temporal | 3.20 (.13) | 3.27 (.13) | 6.877 | .010 | .075 |
| L sup frontal | 3.30 (.15) | 3.35 (.14) | 2.318 | .132 | .027 |
| R sup frontal | 3.22 (.18) | 3.30 (.15) | 4.845 | .030 | .054 |
| R caudal middle frontal | 2.86 (.17) | 2.94 (.12) | 5.140 | .026 | .057 |
| L cuneus | 2.32 (.26) | 2.22 (.22) | 3.699 | .058 | .042 |
| R cuneus | 2.35 (.24) | 2.23 (.20) | 6.957 | .010 | .076 |
| Surface Area | | | *F* | *p* | $\eta_p^2$ |
| Total L hemisphere | 82097.65 (7921.16) | 81644.73 (7239.12) | 0.095 | .758 | .001 |
| Total R hemisphere | 82266.15 (8009.99) | 81864.22 (7421.82) | 0.068 | .796 | .001 |
| L fusiform | 3083.79 (400.62) | 3292.35 (390.85) | 6.779 | .011 | .074 |
| R fusiform | 2973.06 (356.88) | 3220.38 (400.02) | 11.430 | .001 | .119 |
| L lateral occipital | 4981.50 (641.81) | 4699.46 (472.21) | 6.175 | .015 | .068 |
| R superior frontal | 7070.23 (895.38) | 6736.43 (749.70) | 4.155 | .045 | .047 |
| R caudal middle frontal | 2217.31 (398.32) | 2041.84 (365.52) | 5.016 | .028 | .056 |
| Sulcal Depth | | | *F* | *p* | $\eta_p^2$ |
| L superior parietal | -.0015 (.055) | .0317 (.033) | 10.661 | .002 | .111 |
| R superior parietal | -.0521 (.053) | -.0070 (.041) | 18.075 | <.001 | .175 |
| L inferior parietal | .0953 (.046) | .0880 (.039) | .589 | .445 | .007 |
| R inferior parietal | .0880 (.037) | .0787 (.034) | 1.353 | .248 | .016 |
| L supramarginal | .0512 (.050) | .0269 (.032) | 6.845 | .011 | .075 |
| R supramarginal | .1269 (.059) | .0987 (.048) | 5.800 | .018 | .064 |
| L middle temporal | -.1056 (.064) | -.1480 (.056) | 11.486 | .001 | .119 |
| R middle temporal | -.1070 (.067) | -.1548 (.045) | 14.339 | <.001 | .144 |
| L inferior temporal | -.1154 (.050) | -.0646 (.054) | 20.748 | <.001 | .196 |
| R inferior temporal | -.1225 (.054) | -.0756 (.049) | 17.547 | <.001 | .171 |
| L lateral orbitofrontal | .0057 (.051) | .0272 (.047) | 3.983 | .049 | .045 |
| R lateral orbitofrontal | .0286 (.050) | .0494 (.043) | 4.200 | .044 | .047 |
| R cuneus | -.3297 (.073) | -.3622 (.071) | 4.317 | .041 | .048 |
| L precuneus | .1782 (.055) | .1550 (.041) | 5.003 | .028 | .056 |
| R precuneus | .2115 (.056) | .1724 (.049) | 12.168 | .001 | .125 |

*Note:* Group comparisons controlling for sex and age at scan. ROI values extracted from the FreeSurfer Desikan parcellation; M: mean; sd: standard deviation; R: Right; L: Left, inf: inferior, sup: superior, thickness and sulcal depth measured in mm, area measured as mm<sup>2</sup>

Follow-up analyses were conducted to determine if gestational age at birth in the VPT group was correlated with the gray matter metrics of interest. Partial correlations were calculated using values extracted from regions of interest (ROI) from the FreeSurfer automatic parcellation based on areas with significant group differences on the voxel-wise surface maps. After controlling for sex and age at scan, there was a significant partial correlation between GA and thickness in the right middle temporal (*r*[48] = .370, *p* = .008) and left middle temporal (*r*[48] = .314, *p* = .026) regions. For sulcal depth, there were significant partial correlations between GA and the right precuneus (r[48] = −.322, *p* = .022) and right middle temporal (*r*[48] = −.438, *p* =.001) regions. Significant partial correlations between GA and surface area were found in the left fusiform region (*r*[48] = .330, *r* = .019).

## Discussion

We found a number of differences in gray matter cortical morphometry between this group of preschool age children born VPT and children born FT, particularly in cortical thickness. The VPT group had thinner cortex bilaterally within the temporal, superior middle frontal, and parietal/occipital junction and thicker cortex within the dorsal medial occipital cortex bilaterally. The VPT group showed greater sulcal depth in regions of the right medial temporal, and medial partial lobes, and shallower sulcal depth in regions of the bilateral dorsal parietal lobes and ventral inferior frontal lobes compared with the FT group.

Previous studies have demonstrated a similar pattern of thicker cortex in the medial frontal and parietal lobes in children born VPT. These differences were observed from childhood through to adolescence. It may be that in adolescence, brain maturation in children born VPT continues at a slower pace and these areas undergo the regressive processes (e.g., synaptic pruning) typically seen in FT children at younger ages (Mürner-Lavanchy et al., 2014). Therefore, the areas of thicker cortex in our study of 5-year-old VPT children may represent areas where they are lagging behind in this phase of development (i.e., apparent regressive gray matter changes in thickness, and surface area). We replicated some of the previous findings of thinner cortex in the parietal and temporal lobes seen in older children born VLBW (Sripada et al., 2018). These results may indicate that the VPT group is lagging behind in some progressive/proliferative developmental processes that cause gray matter to measure thicker on MRI (e.g., cell production, dendritic arborization, synapse elaboration) that should ordinarily be occurring at this age, based on studies of typical development (Jernigan, Brown, Bartsch, & Dale, 2016). Another possible explanation is that thinner cortex may be related to early “hyper pruning” that occurred during the neonatal period (Zhou et al., 2018). Overall, it is likely that the findings of areas of thinner and areas of thicker cortex in the VPT group compared to the FT group represents interruptions or delays in the typical processes of cell proliferation, dendritic arborization, and/or cell pruning that occur across different brain regions in early childhood. The results from previous studies of older children and adolescents suggest that the neuroanatomical differences we see in this group of 5 year-olds are likely to continue as these children mature.

In contrast to previous studies of older children and young adults, we did not find smaller cortical surface area in in the prefrontal, lateral and ventral temporal, and lateral and medial parietal regions (Lax et al., 2013; Rimol et al., 2016; Sripada et al., 2018). We found that the VPT group had smaller surface area only in the bilateral fusiform region and greater surface area in the right medial frontal cortex and left occipital pole. During typical development, surface area is in the process of expansion during the preschool period (Brown & Jernigan, 2012). These results may indicate that this healthy VPT group is showing the same expansion in cortical surface area as the FT group across most of the cortex, at least at this stage of their development. These discrepancies may also reflect the inclusion of children with a greater number of birth-related risk factors, including lower gestational age at birth in those studies.

This study did not find much overlap in the regions where there were group differences in cortical thickness and surface area. Most early imaging studies of typically developing children have reported initial increases in gray matter thickness followed by thinning later in childhood, but these observations are now in serious doubt given improvements in measuring cortical thickness. Our data are consistent with the observations of Walhovd and colleagues (Walhovd, Fjell, Giedd, Dale, & Brown, 2017) who found that cortical thickness decreases from the earliest ages that were studied (i.e., starting age 3) through early adulthood, including the age range we studied here. Our group of VPT preschoolers were generally similar to the FT group in terms of surface area and sulcal depth measurements. Additional studies, particularly using longitudinal data, would likely be helpful in elucidating these results by measuring the trajectories of cortical morphometry as these children mature (Rimol et al., 2016; Sripada et al., 2018).

Increased gray matter in the occipital lobe has been demonstrated previously in older children born VPT (Lean et al., 2017). Early exposure to visual stimuli because of early exit from the womb, among other factors, puts preterm children at increased risk for visual problems, and associated effects can be seen in the complex network of cortical and subcortical areas associated with visual development (Ramenghi et al., 2010). All children in the study were screened and all were found to have grossly normal vision. However, it is possible that group differences in occipital cortex growth are related to these early differences in visual experiences, as well as the impact of early periventricular disruption.

Additional differences between the groups were found in sulcal depth and cortical thickness in the right middle temporal regions. These measures were also significantly correlated with GA at birth. This finding has been demonstrated in young adults born VPT (Bjuland, Lohaugen, Martinussen, & Skranes, 2013). This area may be vulnerable to damage in the third trimester, and thus structural differences when children are older may correspond to early damage or developmental difference within this region related to preterm birth (Nosarti et al., 2008).

The current study does have some limitations. The children included in this study were recruited to be “low-risk”, without significant neonatal brain injury. Therefore, these results may not generalize to a VPT-born population with more serious brain injury or more serious health complications related to premature birth. Also, these children have a relatively restricted range of GA at birth, with a mean of 29.5 weeks. More significant effects on gray matter morphometry may be present in children with lower GA, particularly those born at < 28 weeks. An additional limitation is the cross-sectional nature of this study. The current analyses cannot answer the question of developmental trajectory and the changes that occur as a child develops in each of the metrics included in this study.

In summary, the results of this study indicated differences in cortical morphometry between preschool-age children who were born VPT and children born FT. At age 5, the most widespread differences were found in measures of cortical thickness. Measures of surface area and sulcal depth were also different between VPT and FT groups. Given the limited research at this age, these results may help elucidate the pattern of brain development in this preterm population from early infancy to the outcomes reported in older children and young adults.

## Acknowledgements

This work was supported by grants R01HD075865 and R01HD061414 from the *Eunice Kennedy Shriver* National Institute of Child Health and Human Development. We would like to thank all the children and their families who participated in this study; Drs. Martha Fuller, Yvonne Vaucher, Terry Jernigan, Joan Stiles, and Anders Dale for their collaboration on this research project; and Drs. Sarah Mattson, and Carrie McDonald for consultation. The authors declare no conflicts of interest.

